# Modulating cellular cytotoxicity and phototoxicity of fluorescent organic salts through counterion pairing

**DOI:** 10.1101/743716

**Authors:** Deanna Broadwater, Matthew Bates, Mayank Jayaram, Margaret Young, Jianzhou He, Austin L. Raithel, Thomas W. Hamann, Wei Zhang, Babak Borhan, Richard R. Lunt, Sophia Y. Lunt

## Abstract

Light-activated theranostics offer promising opportunities for disease diagnosis, image-guided surgery, and site-specific personalized therapy. However, current fluorescent dyes are limited by low brightness, high cytotoxicity, poor tissue penetration, and unwanted side effects. To overcome these limitations, we demonstrate a platform for optoelectronic tuning, which allows independent control of the optical properties from the electronic properties of fluorescent organic salts. This is achieved through cation-anion pairing of organic salts that can modulate the frontier molecular orbital without impacting the bandgap. Optoelectronic tuning enables decoupled control over the cytotoxicity and phototoxicity of fluorescent organic salts through selective generation of mitochondrial reactive oxygen species that control cell viability. We show that through counterion pairing, organic salt nanoparticles can be tuned to be either nontoxic for enhanced imaging, or phototoxic for improved photodynamic therapy.

## INTRODUCTION

Fluorescent dyes offer great potential as both diagnostic and therapeutic agents, and the combined application has been termed “theranostics”. These compounds can be used to improve cancer diagnoses, assist with image-guided surgery, and treat tumors by photodynamic therapy (PDT). Ideal theranostic agents localize in tumors and become activated by a specific wavelength of light to either emit a different wavelength of light that can be detected for imaging, or generate reactive species for PDT^1, 2^. PDT provides double selectivity through the use of both the dye and light, with the goal of minimizing side effects from the dye or light alone^3^. To realize the full potential of fluorescent dyes in biomedical applications, it is necessary to increase their brightness and tissue penetration in order to detect and treat deeply-embedded tumors, while also eliminating unwanted side effects.

Fluorescent dyes that absorb and emit in the near-infrared (NIR) range offer several advantages for both PDT and in vivo imaging applications. While visible light (400-650 nm) travels only millimeters in tissues, NIR light (650–1200 nm) can travel centimeters^4^: 810 nm and 980 nm NIR light have been shown to penetrate 3 cm of skin, skull, and brain tissue^5^. Additionally, visible light absorbance by endogenous biological fluorophores such as heme and flavin groups^6^ causes autofluorescence and weak signal intensity. On the other hand, NIR light is minimally absorbed by biological material, drastically reducing background noise and increasing penetrance^7, 8^. FDA-approved NIR-responsive fluorescent dyes including indocyanine green, 5-aminolevulinic acid, and methylene blue are available and used in medical diagnostics^9^, but are limited due to their low level of brightness. Other commercially available NIR-responsive fluorescent dyes include heptamethine cyanine (Cy7), Alexa Fluor 750, and heptamethine dye IR-808^10–12^. However, these dyes display low brightness, high toxicity, and poor aqueous stability^13^. Recent PDT-based nanocrystals show energy level tunability via surface ligand modification but have poor biocompatibility due to heavy elements and minute absorbance in the NIR range that stem from a lack of oscillator strength near their bandgap. For example, semiconductor nanocrystals have absorption coefficients of ∼10^3^/cm for PbS and PbSe compared to ∼10^6^/cm for cyanines with bandgaps around 850nm – this translates to 1000 times less absorption per nanometer of material by nanocrystals (**Supplementary Figure S1**).

Fluorescent organic salts, composed of a fluorescent ion and a counterion, have been developed to increase aqueous solubility and photostability^14, 15^. The counterion has largely been thought to have little impact on the properties of the fluorescent organic salts. Only a few reports have investigated the impact of the counterion, but have been limited to encapsulated matrices for modestly increasing the quantum yield^16–19^, or have shown no impact on toxicity ^20, 21^. The latter study investigated two anions with a visible rhodamine dye, but showed no significant difference in cell viability between the two key anions in a range of cell lines (Hs578Bst, Hs578T, and MDA-MB-231) and did not investigate phototoxicity^20^. Here, we focus on NIR-responsive polymethine cyanine dyes, which have been used as effective theranostic agents^10, 22^. Heterocyclic polymethine cyanine dyes have been found to preferentially accumulate in tumors and circulating cancer cells^23^ even in the absence of bioconjugation to tumor-targeting molecules. This is thought to occur through a mechanism mediated by increased expression of organic anion transporter peptides (OATPs) and hypoxia-inducible factor 1-alpha (HIF1α), both of which are upregulated in cancer cells. HIF1α promotes tumor angiogenesis and expression of OATPs, which facilitate the uptake of polymethine cyanine dyes^25^, as shown by competitive inhibition of OATP1B3^22^. Lipophilic photosensitizers may also associate into circulating low-density lipoproteins (LDLs) and be imported by cells via ATP-mediated endocytosis^26^. Charged molecules taken up by the cell accumulate in negatively charged organelles such as mitochondria and lysosomes, where light irradiation can induce generation of reactive oxygen species (ROS)^27^. While the exact mode of uptake and localization varies depending on the chemical characteristics of any given photosensitizer, these mechanisms are uniquely active in tumor cells, leading to tumor-specific accumulation and retention^28^.

Cellular toxicity of fluorescent molecules is caused by the combination of: 1) cytotoxicity – toxicity in the dark, independent of photoexcitation; and 2) phototoxicity – toxicity with light illumination, or photoexcitation. While the tumor-specific accumulation of polymethine cyanine dyes reduces their nonspecific toxicity, low levels of systemic toxicity remain due to the cytotoxicity of unexcited molecules^28^. For applications in tumor imaging, both cytotoxicity and phototoxicity need to be eliminated to minimize side effects. For applications in PDT, cytotoxicity must be eliminated, while phototoxicity should be enhanced to selectively kill cancer cells with targeted light therapy.

We recently reported that a range of weakly coordinating anions can modulate frontier molecular orbital levels of a photoactive heptamethine cyanine cation (Cy^+^) in solar cells without changing the bandgap^29, 30^. Thus, we are able to control the electronics (i.e. frontier molecular orbitals) of photoactive molecules independently from their optical properties (i.e. bandgaps). We have subsequently employed this electronic tunability to demonstrate cyanine-based organic salt photovoltaics with > 7 year lifetime under typical solar illumination^31^.

Here, we demonstrate the impact of the counterion on independently controlling both cytotoxicity and phototoxicity of fluorescent organic salts in cancer cells for enhanced imaging and improved PDT (**Figure 1A**). We achieve this by pairing the NIR-absorbing Cy^+^ with various dipole-modulating counterions, and characterizing their effect on human lung carcinoma and metastatic human melanoma cell lines. We find that counterion pairings with small hard anions lead to high cytotoxicity even at low concentrations. In comparison, counterion pairings with bulkier, halogenated anions can remain nontoxic even at 20x higher concentrations. We further report a distinct intermediate group of anion pairings that are highly phototoxic, but exhibit negligible cytotoxicity, making them ideal photosensitizers for PDT. This concept of tuning the cytotoxicity and phototoxicity of fluorescent organic salts is a new platform for controlling the photoexcited interactions at the cellular level. It opens new opportunities for greater tissue penetration and the potential for minimizing side effects. Moreover, this approach may be applied to both novel and existing luminophores, including assembled fluorescent probes, phosphors, nanocrystals, and other hybrid nanoparticles^32–36^.

**Figure 1.**
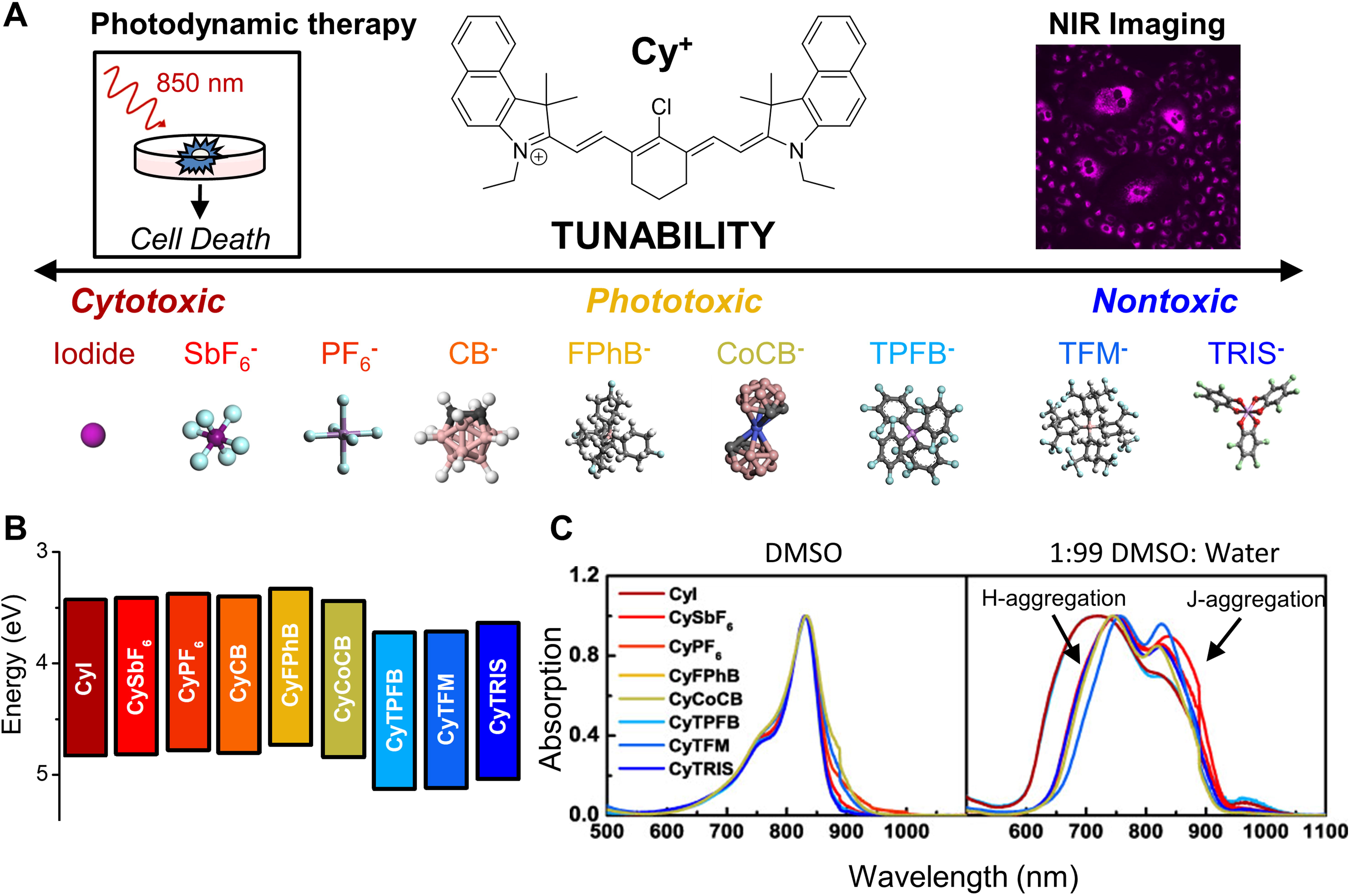
Pairing a fluorescent cation such as heptamethine cyanine (Cy^+^) with varying counterions enables tunability in cellular toxicity through optoelectronic control to improve near-infrared (NIR) imaging and photodynamic therapy. (A) Anions on the left are generally cytotoxic, anions in the middle are selectively phototoxic and ideal for applications in photodynamic therapy, and anions on the right reduce toxicity for applications in fluorescence imaging. Anions: Iodide; hexafluoroantimonate (SbF_6_^−^); hexafluorophosphate (PF_6_^−^); o-carborane (CB^-^); tetrakis(4-fluorophenyl)borate (FPhB^−^); cobalticarborane (CoCB^−^); tetrakis (pentafluorophenyl) borate (TPFB^−^); tetrakis[3,5-bis(trifluoro methyl)]borate (TFM^−^); Δ-tris(tetrachloro-1,2-benzene diolato) phosphate(V) (TRIS). (B) The counterion shifts the HOMO energy level while allowing the interference band gap to remain the same. Ultraviolet photoelectron spectroscopy (UPS) was used to measure the frontier energy levels of Cy^+^ with indicated counterion pairings in the solid state. Reproduced and modified with permission from Suddard et al^28^. (C) Fluorescent organic salts aggregate in aqueous environments. Organic salts fully dissolved in DMSO have a clear maxima at 830 nm with a leading shoulder when characterized with UV-vis spectroscopy. However, in aqueous solution combinations of H- and J-aggregation of organic salts can be seen by blue-shifted peaks (lower wavelength) and red-shifted peaks (higher wavelength), respectively.

## RESULTS

### Characterization of fluorescent organic salts

Cy^+^ is a photoactive cation that absorbs and emits in NIR wavelengths, with a bandgap of 1.3 eV (**Figure 1B**). A range of anions were tested based on our previous studies, demonstrating a full range of valence energy levels that can be tailored by over 1 eV^29, 31^. These include: hard anions iodide (I^-^), hexafluoroantimonate (SbF_6_^−^), and hexafluorophosphate (PF_6_^−^); and bulkier halogenated anions tetrakis(4-fluorophenyl)borate (FPhB^-^), o-carborane (CB^-^), cobalticarborane (CoCB^-^), tetrakis (pentafluorophenyl) borate (TPFB^-^), tetrakis[3,5-bis(trifluoro methyl)]borate (TFM^-^), and Δ – tris(tetrachloro-1,2-benzene diolato) phosphate(V) (referred to as Δ-TRISPHAT, further abbreviated as TRIS^−^) (**Supplementary Figure S2**). The counterion causes distinct shifts in the highest occupied molecular orbital (HOMO) energy levels of heptamethine cyanine salts without changing the size of the bandgap in the solid state (**Figure 1B**). These changes to energy level are found to be consistent for salt nanorparticles in aqueous solution by measuring shifts to the redox potential and zeta potential (**Supplementary Figure S3A-B, Supplementary Table S1**), both of which have been correlated to HOMO^37, 38^. The optical properties of the different ion-counterion pairings remain the same, with equivalent quantum yields and absorbance/emission spectra (**Figure 1C, Supplementary Figure S4, Supplementary Table S2)**. In DMSO, fully dissolved salts display a major peak at 833 nm and a minor shoulder at 764 nm (and no observable shift in redox potential as shown in **Supplementary Figure S3C-D**). Organic salt nanoparticles were formed by diluting these solutions in mixtures of DMSO:H_2_O. All of the organic salts formed soluble nanoparticles with this approach, which is expected due to their similar solubilities in water (**Figure 1C, Supplementary Table S3**). In aqueous solution, the nanoparticles exhibit distinct peak broadening from the major peak and the minor shoulder. The hypsochromic shift of the 764 nm shoulder peak and a bathochromatic shift of the 833 nm peak are indicative of both H- and J-aggregation during the nanoparticle formation process. Nanoparticle organization limits the availability for exchange of the ions and preserves salt composition. Nanoparticle size of a typical bulky pairing (CyTPFB) was characterized by small-angle X-ray scattering (SAXS): the mean particle size is 4.1 ± 0.6 nm (**Supplementary Figure S5A**), a size that is easily taken up by cells^39^. This data was corroborated using scanning electron microscopy (SEM) (**Supplementary Figure S5B**). The nanoparticles remain stable in cell media for at least 22 days (**Supplementary Figure S6**).

### Tunable cellular toxicity

Human lung carcinoma (A549) and metastatic human melanoma (WM1158) cell lines were used as representative models of two distinct cancer types with increased expression of OATP1B1 and OATP1B3^40, 41^ but have limited treatment options. Cells were treated with multiple Cy^+^-anion pairings by diluting organic salts with cell media to generate self-forming nanoparticles. Cells were incubated with various concentrations of the salt nanoparticles with or without 850 nm light to assess cytotoxicity in the dark and phototoxicity with 850 nm irradiation. Cell viability assays show that Cy^+^ is cytotoxic at 1 μM for A549 cells even without exposure to NIR light when paired with small hard anions such as I^-^, SbF_6_^-^, or PF_6_^−^, and only slightly more phototoxic when exposed to light (**Figure 2A**). In contrast, pairings with anions such as FPhB^-^ and CoCB^-^ have little cytotoxicity for concentrations below 7.5 µM but are already highly phototoxic at 5.0 µM and 5.5 µM, respectively (**Figure 2B**). The combination of low cytotoxicity and high phototoxicity is ideal for photosensitizers in PDT. This starkly contrasts to reports that the anion has no impact on dark cytotoxicity in breast cancer cells when paired with a larger bandgap fluorophore^20^. On the other hand, Cy^+^ is found to be nontoxic when paired with TPFB^-^, TFM^-^, and TRIS^−^. These pairings display negligible cytotoxicity and only modest phototoxicity at much higher concentrations, > 15 μM (TPFB), > 20 μM (TFM), and > 30 μ (TRIS), making them more ideal for imaging applications (**Fig 2C**). Both cytotoxicity and phototoxicity are shown to be dose-dependent for all ion pairings tested, with the exception of TRIS^-^, which displayed no cytotoxicity in the concentrations tested up to 100 μM (**Figure 2C**; **Supplementary Table S4**). The dose-dependent response observed in A549 cells is also consistent in WM1158 cells (**Supplementary Figure S7).**

**Figure 2.**
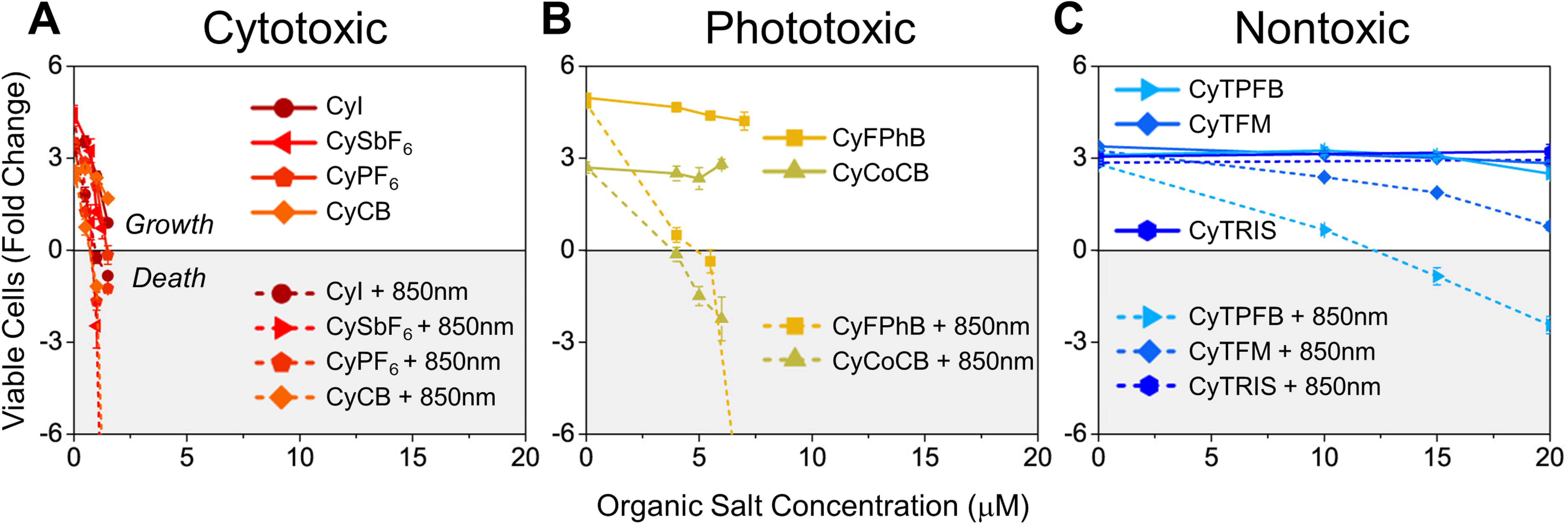
Organic salts with tunable toxicity can be used to target human cancer cells. Toxicity of photoactive cation heptamethine cyanine (Cy^+^) is tuned with anion pairing. Human lung cancer A549 cells were incubated with various concentrations of Cy^+^ with different anionic pairings with or without NIR (850 nm) excitation. Cell viability was determined on day 4 by trypan blue staining and cell counting. (**A**) In A549 cells, CyI, CySbF_6_, CyPF_6_, and CyCB (red/orange) are toxic at low concentrations (1 μM), and cell death occurs independent of light excitation (cytotoxic). (**B**) CyFPhB and Cy B (yellow/green) do not display significant toxicity without light activation, but when photoexcited they induce significant cell death (phototoxic). (**C**) CyTPFB, CyTFM, and CyTRIS (blue) display low toxicity with and without light (nontoxic). Data are displayed as means ± SEM, *n* = 3.

### Mechanism of toxicity

To determine the mechanism of the observed tunability in cytotoxicity and phototoxicity, we investigated salt localization within the cell, which can influence the types of ROS generated and their impact on the cell. Colocalization analysis was done in A549 cells dosed with CyPF_6_ (**Figure 3A**) and stained with a DNA stain, Hoechst (Ho; **Figure 3B**), and a mitochondrial stain, Rhodamine 123 (Rho123; **Figure 3C**). Colocalization was observed for CyPF_6_ and mitochondrial tracker Rho123, but not with DNA-specific Ho (**Figure 3D**). This indicates that the salts preferentially localize in the mitochondria, which is expected due to the positive charge of Cy^+^. Some of the salts that does not colocalize with Rho123 could potentially be found within lysosomes, another negatively charged organelle within the cell. Similar results were observed with CyTPFB and CyFPhB (**Supplementary Table S5**).

**Figure 3.**
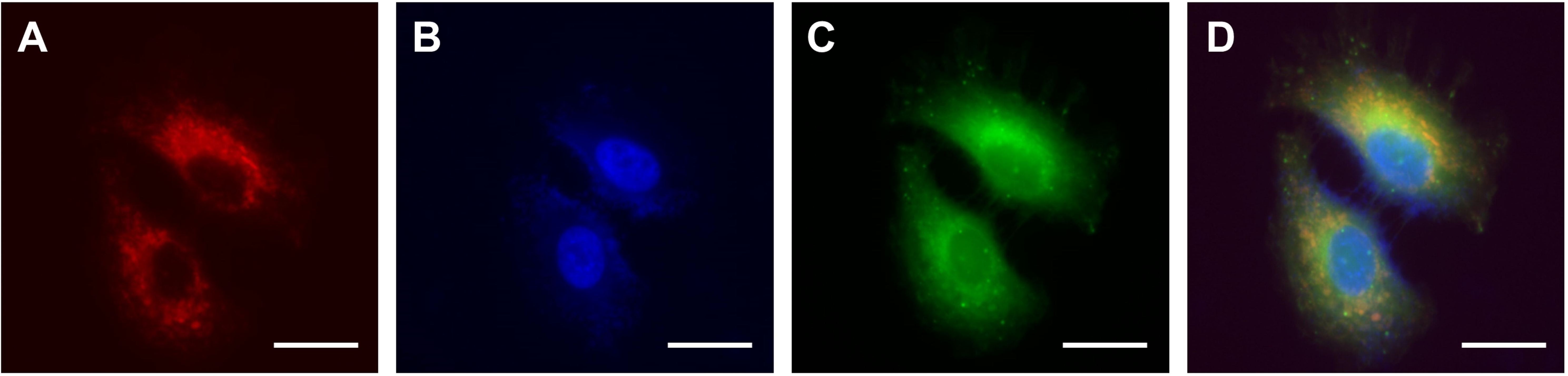
Fluorescent organic salts preferentially accumulate in the mitochondria and lysosomes of cells. A549 cells were treated with 1 μM CyPF_6_. (**A**) CyPF staining. (**B**) DNA lysosomes of cells. A549 cells were treated with 1 μ staining using 2’-[4-ethoxyphenyl]-5-[4-methyl-1-piperazinyl]-2,5’-bi-1H-benzimidazole trihydrochloride trihydrate (Hoechst). (**C**) Mitochondrial staining using Rhodamine 123 (Rho123). (**D**) Superimposed CyPF_6_ + Hoechst + Rho123 staining. Scale bar = 20 µm (100x).

The mechanism of tunability was further studied by oxidative stress analysis using ROS sensitive probes. MitoSOX was used to measure mitochondrial superoxide, and chloromethyl-2′, 7′-dichlorodihydrofluorescein diacetate (Cm-H_2_DCFDA) was used to analyze general cytoplasmic ROS levels in cells treated with phototoxic levels of organic salts (**Figure 4**). We found that an increase in mitochondrial superoxide is directly correlated with both cytotoxicity and phototoxicity of organic salts. Cytotoxic CyPF_6_ generates superoxide with or without light; phototoxic (but not cytotoxic) CyFPhB photo-generates superoxide only with illumination; and nontoxic CyTPFB generates minimal superoxide even with illumination at high concentrations. No cytoplasmic ROS was detected using general cytoplasmic ROS probe Cm-H_2_DCFDA (**Figure 4**). This data demonstrates that the toxicity of organic salts is caused by localized generation of superoxide within the mitochondria. Mitochondrial superoxide is known to mediate apoptosis through oxidative damage of mitochondrial DNA, hyperpolarization of the mitochondrial membrane potential, and protein modifications leading to the opening of the mitochondrial permeability transition pore^42^. A key difference in cells treated with CyPF_6_ is the presence of mitochondrial ROS, even without light excitation. This is likely due to the stability of nanoparticles: UV-Vis spectroscopy showed that while pairings with small, hard anions (CyI, CySbF_6_, and CyPF_6_,) can form nanoparticles in aqueous solution (**Figure 1C**), nanoparticle aggregates were not observed in cell media containing fetal bovine serum (**Supplementary Figure S8).** Pairings with bulkier halogenated anion pairings formed stable and soluble nanoparticles even in cell media. Lack of nanoparticle formation may lead to cytotoxic species, which are toxic even without light activation because they are more likely to interfere with mitochondrial electron transport chain complexes, a process known to generate ROS. In contrast, stable nanoparticles with average size of < 20 nm are still able to enter the cell^39^, but size limitations likely restrict their ability to directly interact and inhibit protein complexes in the mitochondrial membrane.

**Figure 4.**
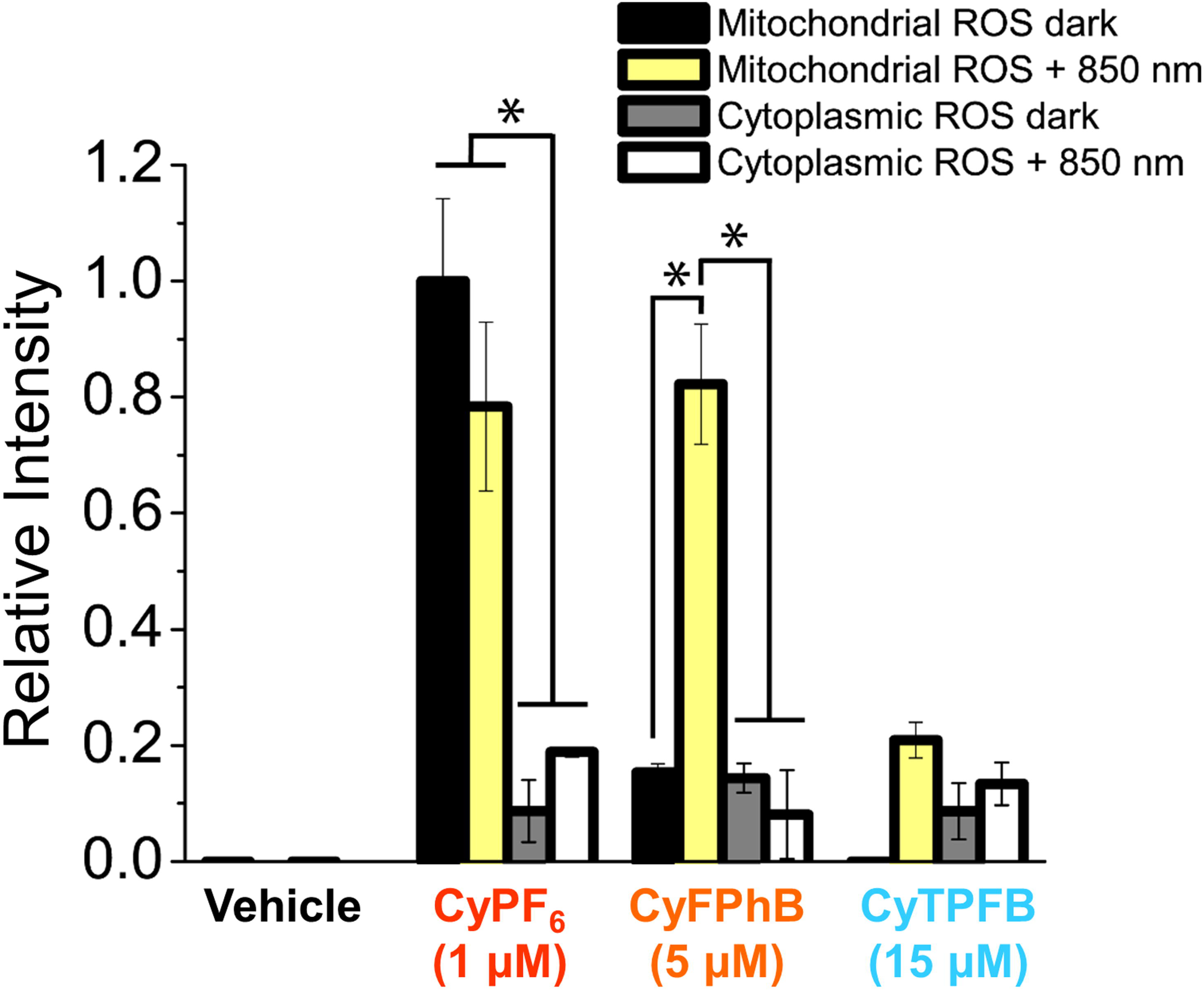
Fluorescent organic salts generate mitochondrial superoxide. MitoSOX was used to measure mitochondrial superoxide, and H_2_DCFDA for general cytoplasmic ROS in A549 cell treated with organic salts at indicated phototoxic concentrations over 4 days. Phototoxic concentrations were determined from the data in Figure 1. This data confirms that CyPF_6_ is cytotoxic, catalyzing superoxide with or without light; CyFPhB is phototoxic but not cytotoxic, photo-generating superoxide only with illumination; and CyTPFB is nontoxic, generating minimal superoxide even with light at high concentrations (\**P* ≤ 0.05). Data are displayed as means ± SD, *n* = 3.

To determine whether the counterion affects cellular uptake of organic salts, intracellular levels of different Cy^+^-anion pairings were measured by high performance liquid chromatography-mass spectrometry (HPLC-MS). No correlation was observed between toxicity and the intracellular concentration of organic salts (**Supplementary Figure S9A**), demonstrating that differential anion mediated uptake is not the cause of the observed modulation in toxicity, even if it is possible that nanoparticle size may be altered upon cellular uptake. In fact, it appears that the opposite may be true: Cy^+^ anion pairings with lower cytotoxicity generally had higher intracellular concentrations. However, it should be noted that toxic salts that induce cell death are more likely to rupture and release dyes, potentially decreasing the observed intracellular concentrations.

Furthermore, we found that the anions themselves are not toxic: addition of a phototoxic anion such as I^-^ paired with a non-fluorescent cation such as potassium (K^+^) is neither cytotoxic nor phototoxic (**Supplementary Figure S9B**). Non-cytotoxic anion-cation pairings cannot be made more toxic by addition of toxic anion salts; for example, a nontoxic salt (CyTRIS) does not become cytotoxic or phototoxic by addition of a toxic precursor salt (KI; **Supplementary Figure S9B**). However, when the reverse experiment was done and a toxic salt (CyPF_6_) was supplemented with a nontoxic precursor salt (KTPFB), toxicity was mitigated (**Supplementary Figure S9C**). This is likely due to variance in nanoparticle stability in cellular environments: in cell media, nanoparticles become less stable when Cy^+^ is paired with small, hard anions (I^-^, SbF_6_, and PF_6_), while nanoparticles remain stable when Cy^+^ is paired with bulkier halogenated anions (FPhB^-^, CoCB^-^, TPFB^-^, TFM^-^, and TRIS^-^; **Supplementary Figure S8**). Thus, CyPF_6_ may undergo an energetically favorable anion exchange with KTPFB to generate the nontoxic CyTPFB species in cell media, leading to decreased toxicity and increased cell viability. These data indicate that the toxicity of organic salts is not due to the toxicity of the anion itself, or cellular uptake.

### Applications in imaging

We next demonstrated that the concept of counterion-mediated tunability can be used to improve *in vitro* imaging of live cells. Commercially available cyanine molecules used for NIR imaging are typically formulated with halide anions (e.g. chloride or iodide), including the Cy3, Cy5, and Cy7 analogs. We performed anion exchange reactions on Cy7Cl to replace the chloride with the range of anions described above. While Cy7Cl is highly cytotoxic, Cy7^+^ can be tuned to become less toxic when paired with TPFB^-^ and TRIS^-^ (**Figure 5A**). This demonstrates that anionic modulation of toxicity is not limited to a specific fluorescent cation, and this effect can be replicated in alternative organic salt formulations. Reduced toxicity is desirable for live cell imaging, as brighter images can be captured with less cellular damage. We have improved live cell imaging using nontoxic anion pairing in both Cy^+^ and Cy7^+^. In contrast to the images obtained using toxic CyPF_6_ and Cy7Cl, brighter images can be captured using nontoxic CyTPFB and Cy7TPFB (**Figure 5B-E**). Due to their high toxicity, CyPF_6_ and Cy7Cl must be used at low concentrations of 1.2 µM and 1.0 µM, respectively. Nontoxic CyTPFB and Cy7TPFB can be used at higher doses of 95 µM and 6 µM, respectively, allowing for increased absorption and absolute brightness while preserving cell viability. Thus, enhanced brightness and lack of toxicity lead to improved images that capture representative cells under less cellular stress.

**Figure 5.**
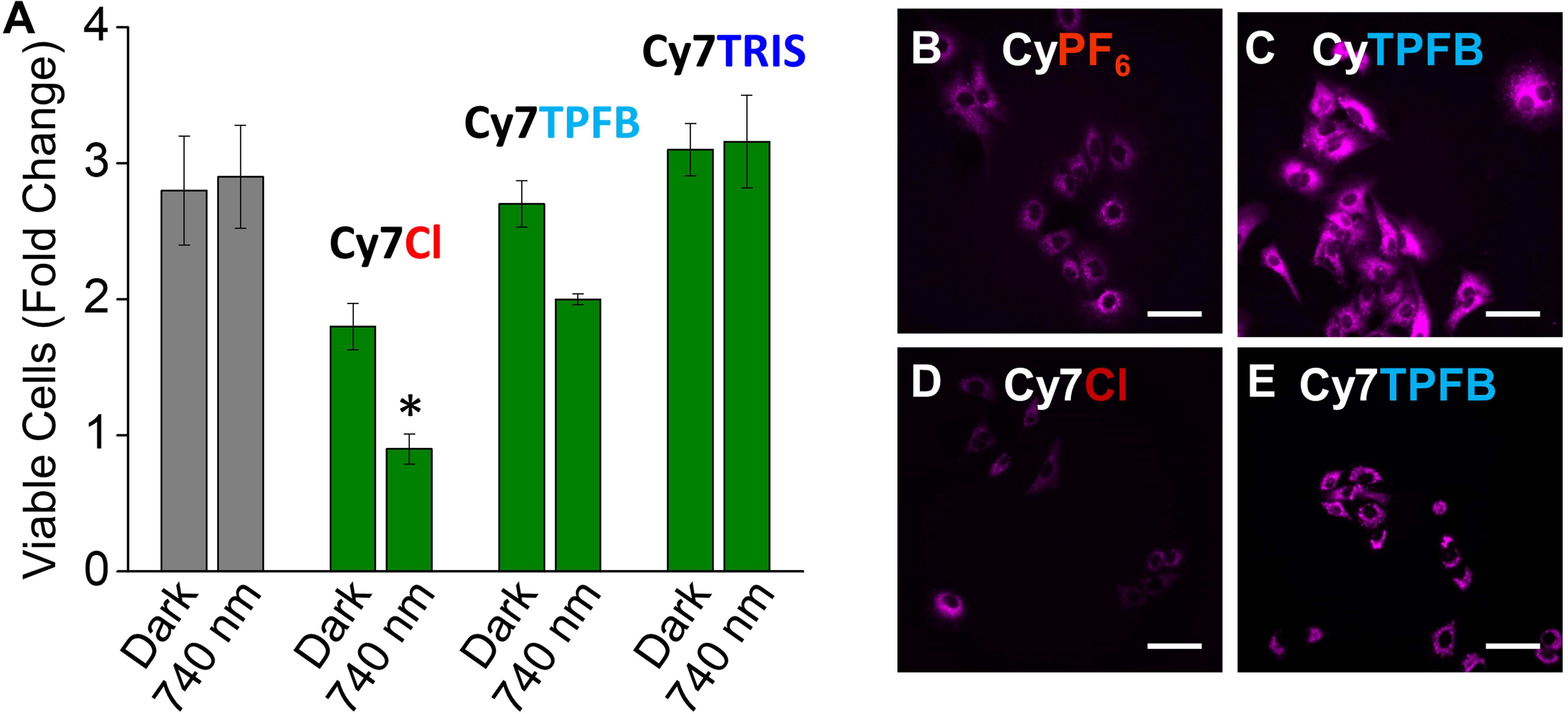
Novel and commercial fluorescent dyes can be tuned to be nontoxic for brighter imaging. (**A**) Commercially available Cy7, sold as Cy7Cl, can be tuned for toxicity through counterion pairing. A549 cells were incubated with Cy7^+^ paired with indicated anions at 1 μM. Commercial formulation of Cy7 with Cl^-^ is found to be cytotoxic; TPFB^-^ pairing shows a dramatic decrease in cytotoxicity with a minor amount of phototoxicity; TRIS^-^ pairing eliminates both cytotoxicity and phototoxicity (\**P* ≤ 0.05). Data are displayed as means ± SEM, *n* = 3. (**B**) Novel fluorescent cation Cy^+^ paired with PF_6_^-^ is cytotoxic at low concentrations (1.2 μM), leading to dim images. (**C**) However, Cy^+^ paired with TPFB^-^ is non-toxic even at increased concentrations (95 μM), and provides brighter images. (**D**) Commercially available Cy7Cl is cytotoxic at 1 μM and provides dim images. (**E**) When Cy7 is paired with counterion TPFB^-^, it also becomes non-toxic at higher concentrations (6 μM) and provides brighter images. Scale bar = 100 µm (40x).

## DISCUSSION

There is growing interest in developing noninvasive cancer theranostic agents that can detect and target a wide range of tumor types with minimal toxic side effects. This work develops a platform for tuning the toxicity of theranostic agents through counterion pairings for applications in both enhanced imaging and effective therapy. We have demonstrated the ability of weakly coordinating anions to tune cellular toxicity of multiple organic salts, by influencing the energy level of the fluorescent cation to impact generation of mitochondrial superoxide. Nanoparticle formation is necessary for the observed modulation of cellular toxicity by the counterion, as it preserves salt composition and prevents ionic dissociation in aqueous solution. We have shown that the tunability in cellular toxicity is independent of intracellular concentration, localization, anionic toxicity, and is not specific to a particular ionic fluorophore. These data demonstrate that electronic modulation via counterion pairing can tune the cytotoxicity and phototoxicity of photosensitizers in cellular environments. We find a correlation between the zeta potential of nanoparticles in aqueous solution and their cyto- and phototoxicity (**Supplementary Figure S10**). Cytotoxic/phototoxic nanoparticles have positive zeta potentials, while non-cytotoxic/phototoxic nanoparticles have negative zeta potentials, and non-cytotoxic/non-phototoxic nanoparticles have even lower negative zeta potentials (**Supplementary Figure S10**). Interestingly, nanoparticles with negative zeta potentials have distinct cyto- and phototoxicities, while nanoparticles with positive zeta potentials have overlapping cyto- and phototoxicities (**Supplementary Figure S10**). Zeta potential has been correlated to HOMO level^38^, and these shifts in HOMO are likely the driving force for dictating phototoxicity. While we have demonstrated the correlation between energy level modulation and phototoxicity, the mechanism by which valence energy levels effect cellular toxicity remains an open question for future studies. We hypothesize that the degree of phototoxicity is dictated by energy level resonance with components in the mitochondria. For example, CyFPhB with a lower absolute HOMO is highly phototoxic, while CyTPFB with a higher absolute HOMO is not phototoxic even at orders of magnitude higher concentrations. This is likely due to the ability of the photoactivated fluorophore to resonately perform electron transfer reactions within the mitochondria and therefore produce varying amounts and types of particular radical and reactive species. Energy level modulation is only achievable with nanoparticle formulation, as free salts show the same redox potential and therefore the same energy level (**Supplementary Figure S3**)^29^. Non-toxic pairing with anions such as TPFB^-^ and TFM^-^ can be used to reduce cellular toxicity during diagnostic imaging. In contrast, we have selectively enhanced phototoxicity in response to NIR excitation while eliminating dark cytotoxicity of Cy^+^ across a range of cell lines by pairings with anions such as FPhB^−^ and CoCB^−^. This approach has the potential to increase targeting efficacy in tumors while minimizing nonspecific toxicity in healthy tissue. In addition to having broad clinical applications, this work gives insight into a novel method for modulating the electronic characteristics of fluorescent cation-anion pairings, and provides a rational strategy for enhancing existing photodynamic drugs and imagers.

## METHODS

### Synthesis

#### Synthesis of CyPF_6_, CySbF_6_, CyFPhB, and CyTPF

Precursor salts (CyI and NaPF_6_, NaSbF_6_, NaFPhB, or KTPFB) were dissolved in methanol:dichloromethane (MeOH:DCM) mixtures and stirred at room temperature under nitrogen. The counterion precursor was added in 100% molar excess to drive the exchange of ions. The product compounds were formed as solid precipitates after approximately 5 minutes. They were collected using vacuum filtration and rinsed with MeOH. The crude product was dissolved in minimal DCM and run through a silica gel plug with DCM as the eluent to remove unreacted precursors and other impurities. The product compound exiting the silica was recognized by its color and collected. Excess DCM was removed in a rotary evaporator. Reaction yield and purity were confirmed using a high mass accuracy time-of-flight mass spectrometer coupled to an ultra high performance liquid chromatography (UPLC-MS) in positive mode to quantify cations, and in negative mode to quantify anions. For ion purity measurements, solutions of precursors and products were prepared in various known concentrations and analyzed by UPLC-MS. Typical reactions led to products yields of > 60% with purities > 95%. Reaction schemes and purification procedures described previously were used^30, 43^.

#### Synthesis of CyTRIS and CyTFM

Precursor salts (CyI and TBA-TRIS or NaTFM) were dissolved in DCM in a 1:2 molar ratio and stirred at room temperature under nitrogen for 1 hour. The reaction contents were passed through a silica gel plug using DCM as the eluent, where the purified product was collected and quantified with UPLC-MS as described for the salts above. Similar yields and purities were achieved for CyTRIS and CyTFM as other salts.

#### Synthesis of CyCoCB

Precursor salts CyI and NaCoCB were dissolved in MeOH in a 1:2 molar ratio and stirred at room temperature under nitrogen. CyCoCB formed and precipitated out of solution after approximately 5 minutes. The crude product was collected using vacuum filtration and rinsed with MeOH. It was then purified with silica gel chromatography and the purity was quantified with UPLC-MS as detailed previously. Reaction yield and purity of CyCoCB was similar to that of the other salts discussed here^30, 31^.

#### Synthesis of Cy7PF_6_, Cy7FPhB, Cy7TPFB, and Cy7TRIS

Precursor salts (Cy7Cl and NaPF_6_, NaFPhB, KTPFB, and TBA-TRIS) were dissolved in DCM in a 1:2 molar ratio and stirred at room temperature under nitrogen for 1 hour. Reaction contents were passed through a silica gel plug using DCM as the eluent, where the purified product was collected and quantified with UHPLC-MS. Reaction yields were 45-50% with similar purity to other salts discussed here.

*Cyanine7 NHS ester (Cy7)* was utilized as received (Lumiprobe), as a commercial reference.

### Cell culture

Human lung carcinoma (A549) and metastatic human melanoma (WM1158) cells were cultured in Dulbecco’s Modified Eagle’s Medium (DMEM) with 4.5 g/L glucose without sodium pyruvate with 10% heat inactivated fetal bovine serum supplemented with 1 mM glutamine and 1% penicillin and streptomycin. Cells were incubated in 37°C with 5% CO_2_ without light exposure.

### Viability studies

A549 and WM1158 cells were seeded at a density of 50,000 cells per well in 6-well tissue culture plates. After 24 hours of incubation, media was aspirated and replaced with media containing fluorescent dyes at indicated concentrations. Each well was irradiated with an 850 nm LED lamp with an illumination flux of 526mW/cm^2^ for an hour in the incubator, and control cells were left in a dark incubator without irradiation. For studies using Cy7, a custom made 740nm LED lamp was used, but with the same illumination flux. Immediately after irradiation, the media was replenished with fresh dye-laced media and allowed to incubate for another 24 hours. The same procedure was done at 48 and 72 hours, but the cells received no further dye-laced media after 72 hours. Viable cell number was determined at 24 and 96 hours using 4% trypan blue and a Nexcelom Cellometer Auto T4 cell counter. All assays were done with 3 biological replicates. The fold change in cell proliferation over days of treatment was calculated using the following equation^44^:

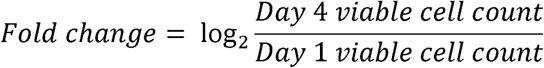

The half maximal inhibitory concentration (IC_50_) was calculated by linear regression analysis of cell viability versus concentration data.

### Fluorescent imaging

Images were obtained using a Leica DMi8 microscope with a PE4000 LED light source, DFC9000GT camera, and LAS X imaging software. A549 cells were seeded in 3 cm tissue culture plates at a density of 50,000 cells per well in DMEM containing fluorescent organic salts at indicated concentrations. The cells were incubated for 2 days at 37°C with 5% CO_2_ until the day of imaging. For live cell imaging, the media was aspirated, and the cells were washed with phosphate buffered saline (PBS, Sigma-Aldrich) 5 times before being imaged in PBS.

For colocalization analysis, A549 cells were grown on 0.5 mm coverslips placed in 3 cm tissue culture plates containing media for 3 days. Cells were then fixed by aspirating media, washing with PBS 5 times, then submerging the coverslip in cold methanol and incubating on ice for 15 minutes. The fixed cells were stained with 1 µM 2’-[4-ethoxyphenyl]-5-[4-methyl-1-piperazinyl]-2,5’-bi-1H-benzimidazole trihydrochloride trihydrate (Hoechst 33342, Invitrogen) for 5 minutes, washed with PBS, and then incubated with 15 µM of 3,6-diamino-9-(2-(methoxycarbonyl)phenyl chloride (Rhodamine123) and 1 µM CyPF_6_ for 15 minutes before being washed and mounted to slides with Fluoromount-G (invitrogen). Cells were analyzed using a Leica DMi8 microscope with a PE4000 LED light source, DFC9000GT camera, and LAS X imaging software.

### Flow cytometry

Cells were incubated with phototoxic concentrations of CyPF_6_ (1 µM), CyFPhB (5 µM), or CyTPFB (15 µM) and exposed to NIR light for 4 days as described above. Each day, cells were collected for analysis by trypsinization from plates (prior to any illumination), spun down and resuspended in a staining buffer consisting of Hank’s buffered salt solution (HBSS, Sigma-Aldrich) with 10 mM 4-(2-Hydroxyethyl)piperazine-1-ethanesulfonic acid (HEPES, Sigma-Aldrich) and 2% FBS. Cells were separated into 2 populations for staining with 15 µM of chloromethyl-2′, 7′-dichlorodihydrofluorescein diacetate (cm-H_2_DCFDA, Invitrogen) for 60 minutes, or 2.25 µM of MitoSOX (Invitrogen) for 20 minutes. Hydrogen peroxide was used as a positive control for H_2_DCFDA. Cells were analyzed on a BD LSR II using FITC and PE-A channels and 30,000 events counted. Fluorescence was normalized to the initial value.

### Ultraviolet visible spectroscopy

Cyanine dyes were diluted to a concentration of 5 µM in cell media. All dyes were characterized using a Perkin-Elmer 25 UV-Vis spectrometer in the wavelength range from 500-1100 nm in normal incidence transmission mode with a resolution of 1 nm and a 1.27 cm path length. A pure solvent reference was utilized to remove reflections so that the absorption is calculated as 1-transmission.

### Zeta potential measurements of nanoparticles

A Zetasizer Nano Z (Malvern Instruments, UK) at 25°C with a 633 nm laser was used to calculate zeta-potential measurements (ζ) using laser Doppler micro-electrophoresis. Measured electrophoretic mobilities (μ*_e_*) were converted to zeta potentials from the Henry equation:

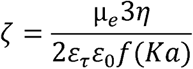

where *ε_τ_* is the dielectric constant of the medium, *ε*_0_ is the permittivity of the vacuum, *f*(*Ka*) is Henry’s function, and *η*, is the viscosity of the colloid. Samples were run in triplicate at a concentration of 10 µM in 10% phosphate buffered saline and 1% DMSO.

### Scanning electron microscopy

Polished glass substrates (Xin Yan Technology LTD) for SEM imaging were cleaned by sonicating in soap, deionized water, acetone, and by boiling in isopropanol for 6 minutes each, followed by oxygen plasma treatment for 3 minutes. Nanoparticles of CyTPFB were spin-coated with 50 µL of 0.5, 5, and 50 µM solutions on a glass substrate at 2000 rpm for 30 seconds. A thin film of tungsten was deposited on the SEM samples to reduce charging. A Carl Zeiss EVO LS 25 Variable Pressure Scanning Electron Microscope was used to capture SEM images of the organic salt nanoparticles. Size distributions were obtained using ImageJ software.

### Small-angle x-ray scattering

Small-angle X-ray scattering (SAXS) was performed with a Rigaku Ultima IV X-ray Diffractometer in the Robert B. Mitchell Electron Microbeam Analysis Lab at the University of Michigan. Parallel beam and SAXS alignment procedures were performed to prepare the diffractometer for measurements. Boron-rich glass capillaries with 1.5 mm outside diameter were purchased from the Charles Supper Company for these measurements. A control sample of 1 mg/mL PbS quantum dots (Millipore Sigma, 3 nm nominal size) in toluene was run first to verify measurement accuracy, producing a mean particle size of 3.3 ± 0.2 nm after subtracting a toluene background scan. A sample of 2.6 mg/mL CyTPFB nanoparticles in 50% DMSO, 50% water was tested, along with a 50% DMSO, 50% water blank, and produced a size distribution curve shown in **Supplementary Figure S5** with a mean particle size of 4.1 ± 0.6 nm. Solubility limits prevented collection of SAXS data for the other CyX nanoparticles, as high concentrations of > 1 mg/mL were necessary to obtain data above the background (see **Supplementary Table S3**).

### Differential pulse voltammetry

Differential pulse voltammetry measurements were made using a μAutoLabIII potentiostat to evaluate oxidation potentials for monomer and nanoparticle salts. For monomers, salts were dissolved at 1 mM in acetonitrile with 100 mM TBA-PF_6_ as a supporting electrolyte. Glassy carbon, Ag/AgNO_3_ (0.36 V vs SCE), and Pt mesh were used as the working, reference, and counter electrodes, respectively. Nanoparticles of the salts were made at 0.1 mM in 10% DMSO (CyI, CyPF_6_) and 50% DMSO (CyFPhB) in H_2_O with 100 mM NaCl as a supporting electrolyte. CyTPFB NPs have greater solubility and were tested at 0.5 mM in 50% DMSO. Ag/AgCl reference electrode (−45 mV vs SCE) was used for the nanoparticle measurements. Nanoparticle solubility limitations in acetonitrile limited collection of CV scans to CyI, CyPF_6_, CyFPhB and CyTPFB, for which concentrations of at least 0.1 mM were achievable.

### Photoluminescence

Photoluminescence (PL) spectra were collected using a PTI Spectrofluorometer for monomers of Cy7X and CyX salts, as well as nanoparticles of CyX salts. For Cy7X monomers, solutions at 1 μM salt in DMSO were used. Solutions of 5 μM CyX salts were prepared in DMSO for PL measurements. Nanoparticles of CyX salts were μM in 1% DMSO, 99% water. A mounted Thorlabs 735 nm LED was used at approximately 5% power as the excitation source for the PL spectra of the CyX monomers and nanoparticles, while a monochromated Xenon lamp (700 nm) was used as the excitation source for Cy7X PL.

### Quantum yield

Quantum yield (QY) data was gathered using a PTI Spectrofluorometer with an integrating sphere (350-900 nm) for monomers of Cy7X and CyX salts. A Thorlabs 735 nm LED was used at approximately 5% power as the excitation source for all quantum yield measurements. Cy7X solutions were made at 1 μM in DMSO, while CyX salts were prepared at 2.5 μM in DMSO.

### Determination of intracellular organic salt concentrations

Cells were seeded at a density of 50,000 cells per well in 6-well plates in media containing 1µM of indicated dye. Cells were allowed to incubate for 3 days at 37°C with 5% CO_2_ with a media change to fresh dye-laced media on day 2. For extraction, media was aspirated from each well, and cells were washed with PBS. The cells were removed from the plate using 0.05% trypsin/EDTA (Thermo Fisher) and centrifuged at 1,500 rpm for 6 minutes. The supernatant was aspirated, and the cell pellet was washed with saline. Saline was aspirated, and the pellets were resuspended with room temperature HPLC-grade 3:7 methanol:acetonitrile (Sigma Aldrich) and centrifuged at 13,000 rpm for 5 minutes. The supernatant was collected in a separate tube, and the pellet was again resuspended in HPLC grade 3:7 methanol:acetonitrile and centrifuged at 13,000 rpm for 10 minutes. The supernatant was combined with the first supernatant for analysis by liquid chromatography-mass spectrometry.

Cell extracts were analyzed the day of extraction using a Waters Xevo G2-XS QToF mass spectrometer coupled to a Waters Acquity UPLC system. The UPLC parameters were as follows: autosampler temperature, 10°C; injection volume, 5 µl; column temperature, 50°C; and flow rate, 300 µl/min. The mobile solvents were Solvent A: 10mM ammonium formate (Sigma-Aldrich) and 0.1% formic acid (Sigma-Aldrich) in 60:40 acetonitrile:water; and Solvent B: 10mM ammonium formate and 0.1% formic acid in 90:10 isopropanol:acetonitrile. Elution from the column was performed over 5 minutes with the following gradient: t = 0 minutes, 5% B; t = 3 minutes, 95% B; t = 4 minutes, 95% B; t = 5 minutes, 5% B. ESI spray voltage was 3,000 V. Nitrogen was used as the sheath gas at 30 psi and as the auxiliary gas at 10 psi, and argon as the collision gas at 1.5 mTorr, with the capillary temperature at 325°C.

Data were acquired and analyzed using MassLynx 4.1 and QuanLynx software. Cy^+^, which typically elutes at 2.5 minutes, was analyzed in positive mode. Standards of each anion-cation pair were run at concentrations of 5, 10, 25, 50, and 100 nM to generate standard curves for quantitation. Blanks were run before each sample to minimize sample carryover.

### Statistical analysis

Statistical analysis was done using OriginPro 8 software. For analyses with more than two group comparisons, a one-way ANOVA analysis was performed with an ad hoc Bonferroni test. To assess the homogeneity of variance and suitability for ANOVA analysis, a Levene’s test was performed. *P*-values < 0.05 are reported as statistically significant.

## ACKNOWLEDGEMENTS

The authors thank Elliot Ensink, Martin Ogrodzinski, Shao Thing Teoh, and Lei Yu for helpful discussions and critical reading of this manuscript. The authors also thank the Michigan State University (MSU) Flow Cytometry Core and the MSU Mass Spectrometry and Metabolomics Core. The authors thank the University of Michigan Robert B. Mitchell Electron Microbeam Analysis Lab.

## AUTHOR CONTRIBUTIONS

S.Y.L. and R.R.L. conceived the project. D.B., M.B., M.Y, R.R.L, and S.Y.L. designed the experiments. D.B. performed all cell toxicity assays as well as mass spectrometry, cell imaging, and flow cytometry experiments. M.B. collected optical and characterization data of the nanoparticles. M.J. assisted with cell toxicity assays. M.B. and M.Y. synthesized the organic salts and nanoparticles. J.H. and W.Z. assisted with zeta potential measurements. A.R. and T.H. assisted with differential pulse voltammetry measurements. B.B. assisted with chemical synthesis. D.B., M.B., R.R.L, and S.Y.L. wrote the manuscript.

## DATA AND MATERIALS AVAILABILITY

All data is available in the main text or the supplementary materials. All data, code, and materials used in the analysis are available upon reasonable request to any researcher for purposes of reproducing or extending the analysis.

