## Supplementary Material for "Modulating cellular cytotoxicity and phototoxicity of fluorescent organic salts through counterion pairing"

Deanna Broadwater^†^, Matthew Bates^†^, Mayank Jayaram, Margaret Young, Jianzhou He, Austin L. Raithel, Thomas W. Hamann, Wei Zhang, Babak Borhan,

Richard R. Lunt^*^, and Sophia Y. Lunt^*^

^†^Equal contribution

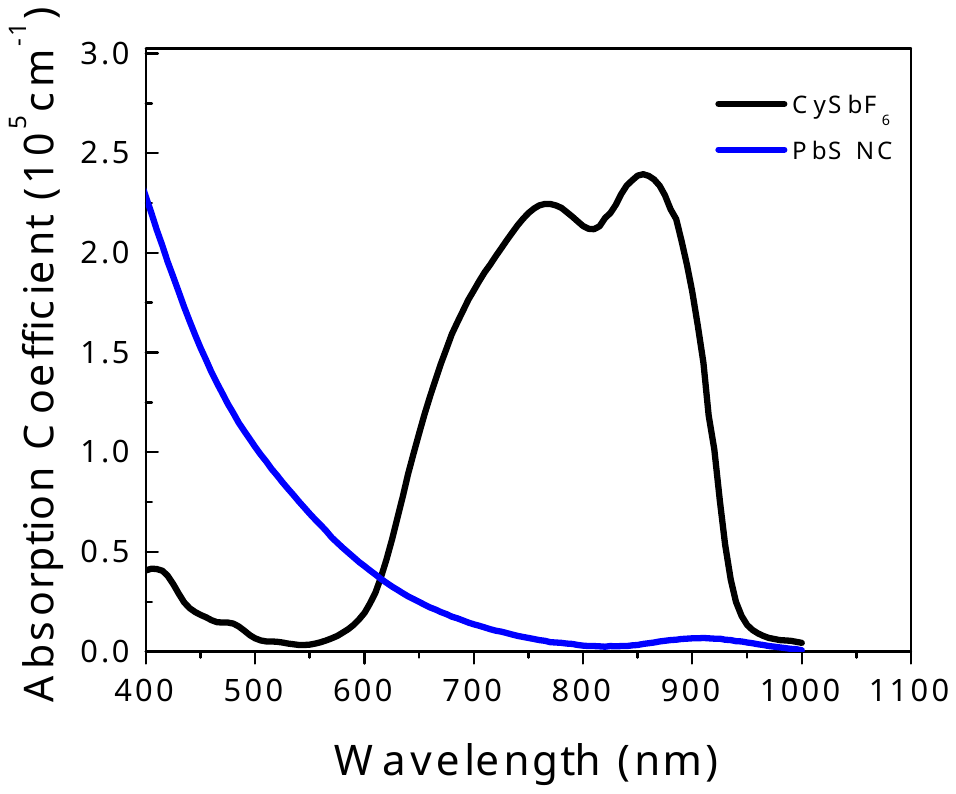

**Supplementary Figure S1.** Solid state absorption coefficient versus wavelength comparison for an exemplary organic salt (CySbF_6_) and nanocrystal (PbS), both with bandgap around 1.3 eV. The organic salt has an absorption coefficient that is orders of magnitude larger than that for the nanocrystal at wavelengths in the near-infrared around the bandgap (650-950 nm).

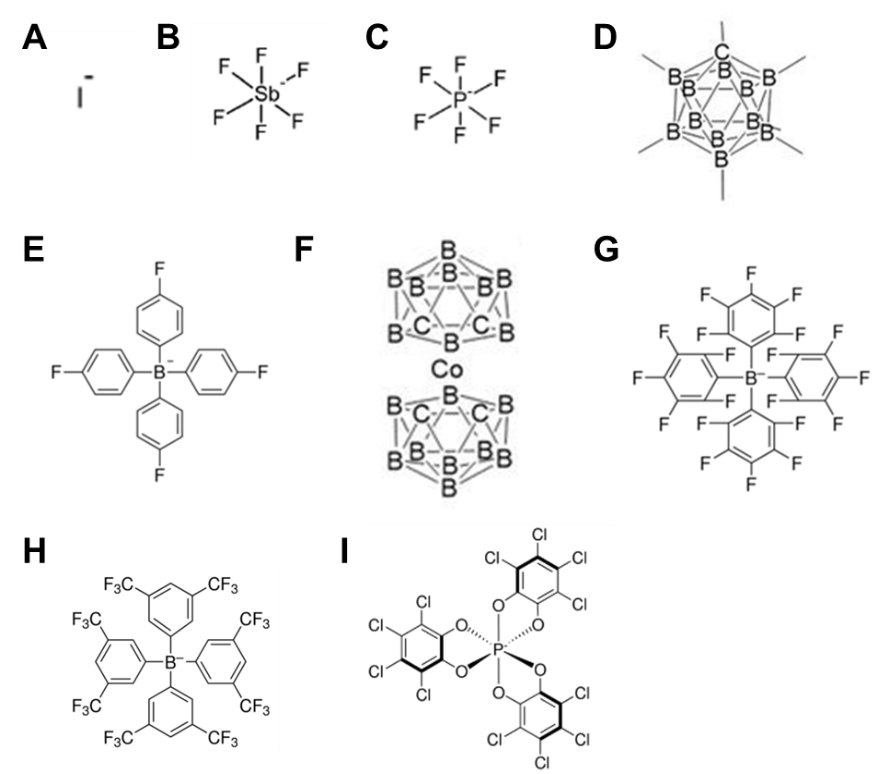

**Supplementary Figure S2.** Molecular structures of anions investigated: (**A**) Iodide; (**B**) hexafluoroantimonate; (**C**) hexafluorophosphate; (**D**) o-carborane; (**E**) tetrakis(4-fluorophenyl)borate; (**F**) cobalticarborane; (**G**) tetrakis (pentafluorophenyl) borate; (**H**) tetrakis[3,5-bis(trifluoro methyl)]borate; and (**I**) Δ-tris(tetrachloro-1,2-benzene diolato) phosphate(V).

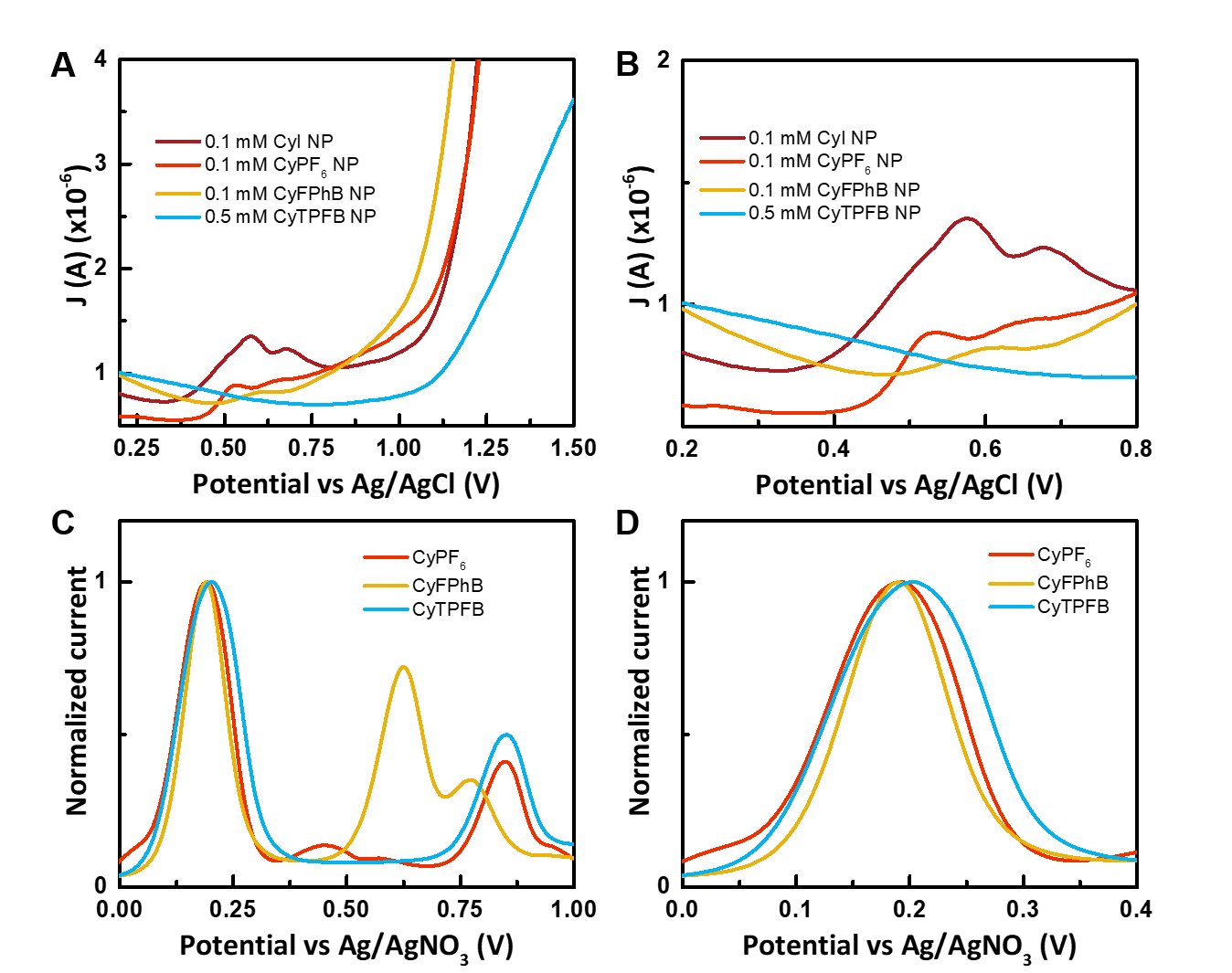

**Supplementary Figure S3.** Differential pulse voltammetry measurements. (**A**, **B**) Nanoparticles of the salts at 0.1 mM in 10% DMSO (CyI, CyPF_6_) and 50% DMSO (CyFPhB) in water. CyTPFB nanoparticles have greater solubility and were tested at 0.5 mM in 50% DMSO. (**C**, **D**) Monomer solutions of a representative cytotoxic, phototoxic, and nontoxic salt in acetonitrile. Monomers demonstrate similar initial oxidation peaks, while nanoparticles have different peak locations. The CyTPFB nanoparticle oxidation peak is outside the redox window available for DMSO:H_2_O mixtures. Anionic effects on the HOMO level measured in the solid state with UPS correlate with redox level shifts: CyPF_6_ shifts to lower potential compared to CyI and CyTPFB is shifted deeper by more the 500mV. Monomers do not show shifts in the first redox peak as their electronic environments are identical after dissociation. DMSO:water solutions were measured with a Ag/AgCl reference electrode (-45 mV vs SCE) and acetonitrile solutions were measured with a Ag/AgNO_3_ electrode (0.36 V vs SCE).

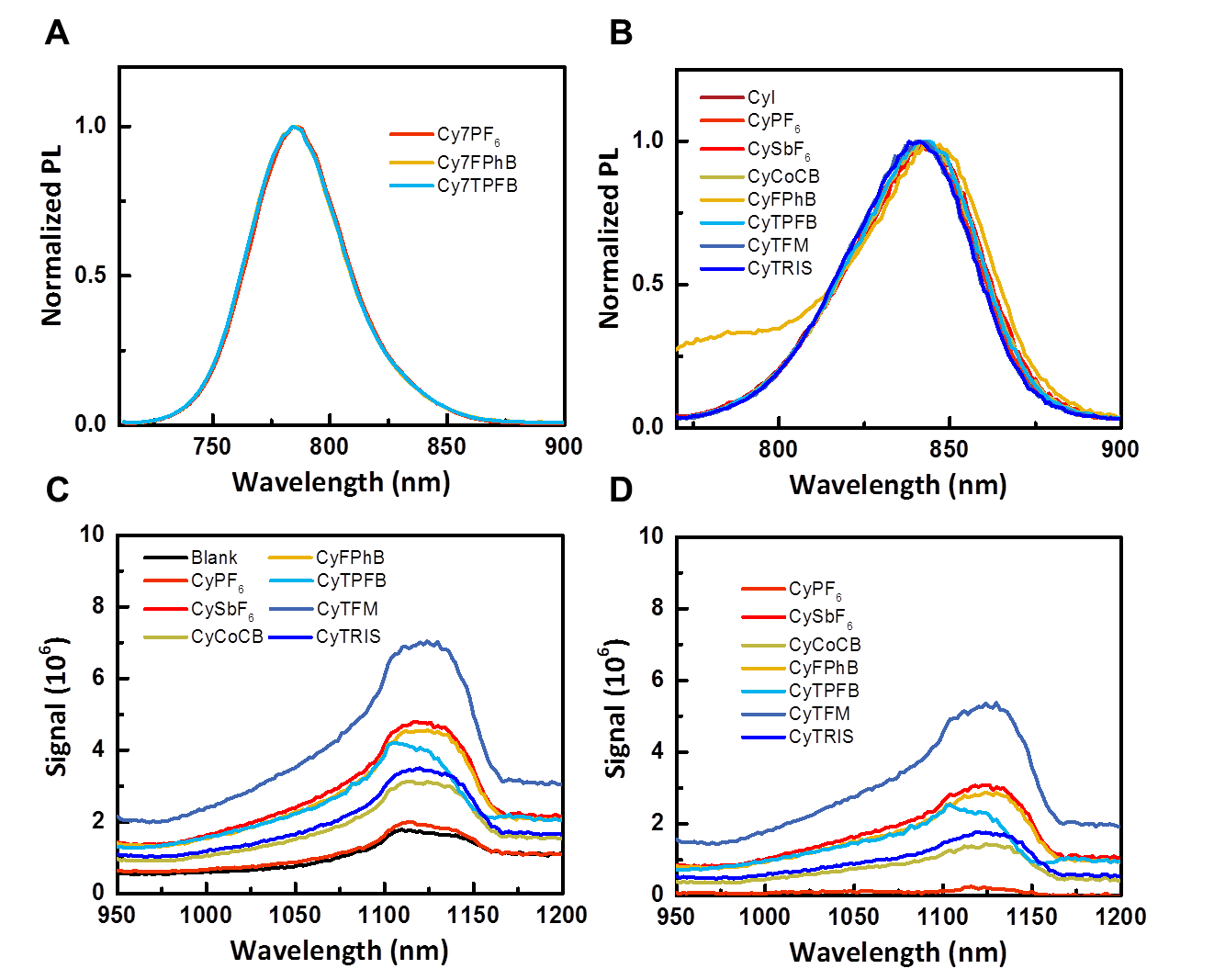

**Supplementary Figure S4**. Photoluminescence (PL) measurements of Cy7X and CyX salts. (**A**) Normalized PL of 1 μM Cy7X monomers in DMSO. (**B**) Normalized PL spectra for 5 μM CyX monomers. (**C**) PL spectra of 2.5 μM CyX nanoparticles compared with the blank (1% DMSO, 99% water). (**D**) Solvent background corrected PL of CyX nanoparticles.

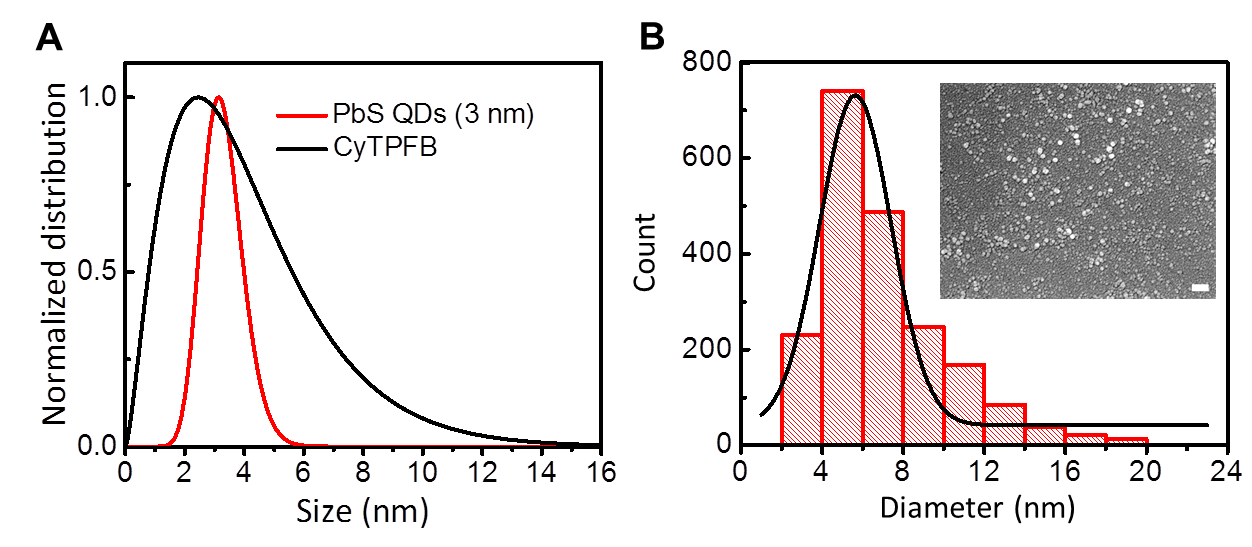

**Supplementary Figure S5.** Nanoparticle aggregation size distribution measurements. (**A**) SAXS measurements of CyTPFB nanoparticles. Mean particle size is 4.1 ± 0.6 nm. PbS quantum dot size distribution is shown as a control with nominal size of 3 nm. (**B**) SEM images (inset, scale bar = 100 nm) of CyTPFB nanoparticles. Mean aggregate size is 7 ± 3 nm with no observable precipitation.

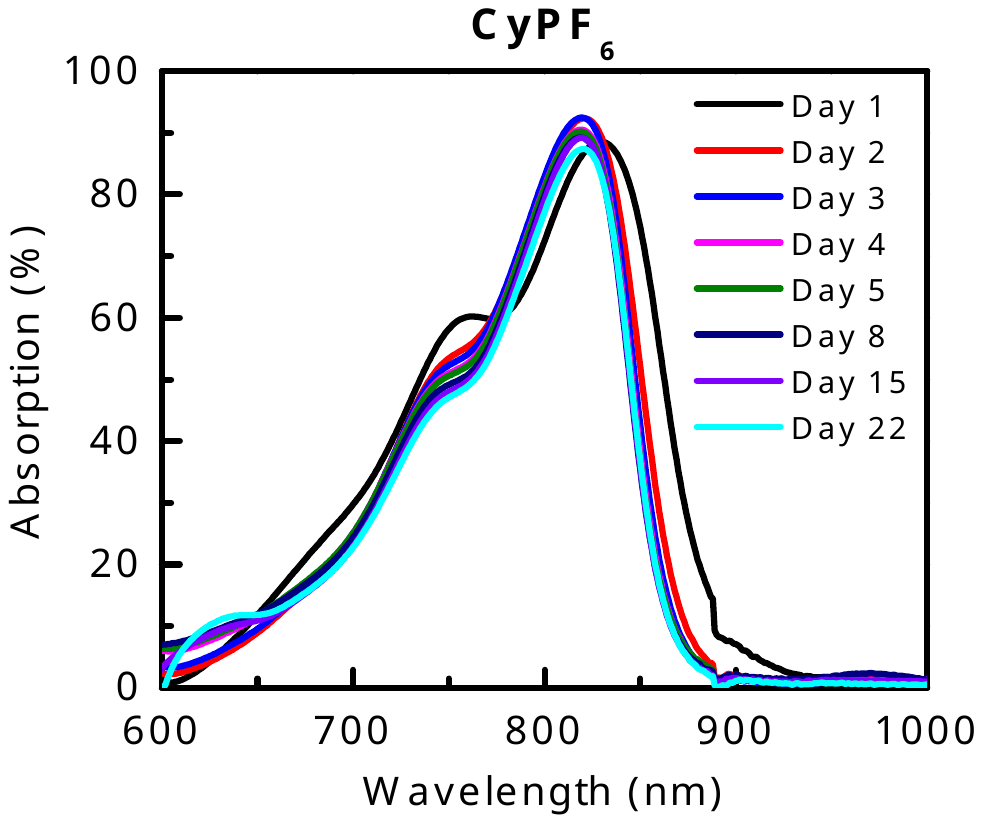

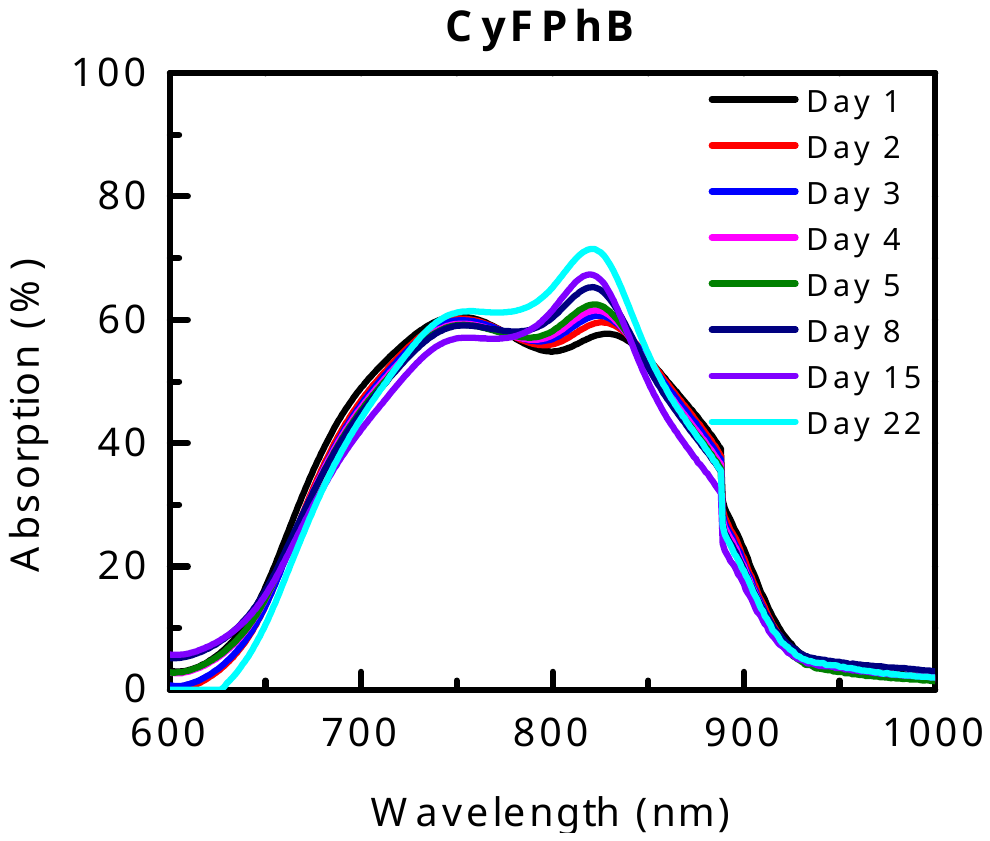

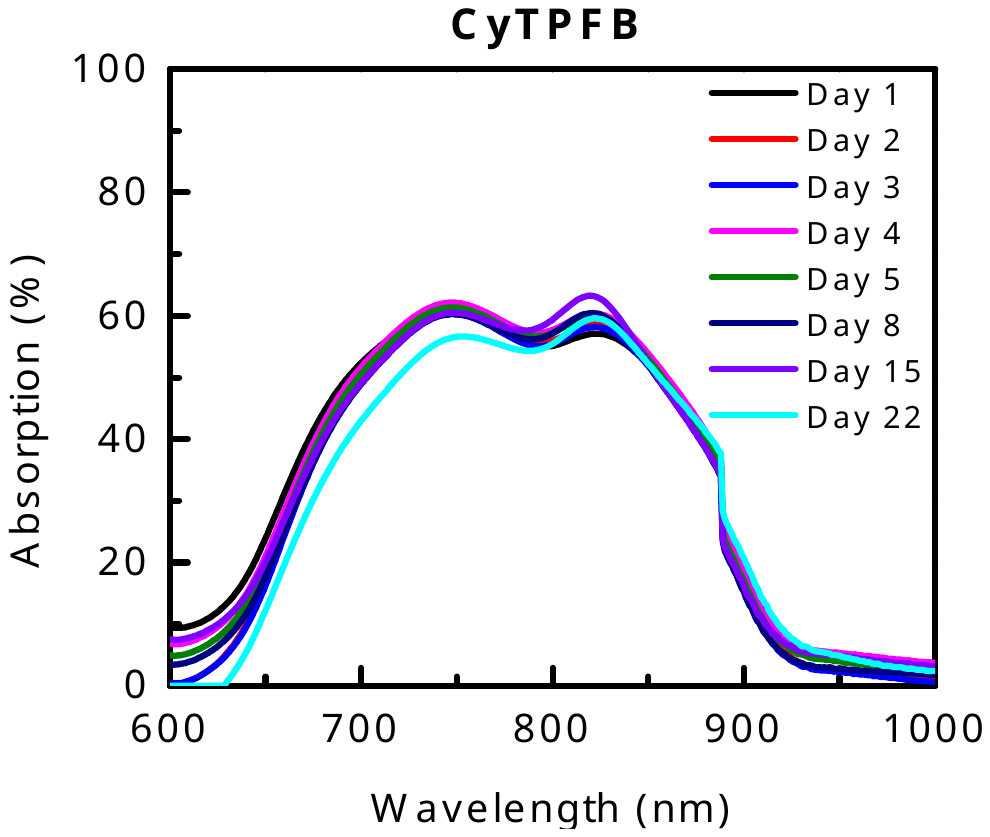

**Supplementary Figure S6.** Nanoparticle lifetime and stability. CyPF_6_ does not form nanoparticles in cell media but nonetheless demonstrates a stable chromophore. Lifetime absorption (100-%T) data collected with UV-Vis spectroscopy for 5 µM CyPF_6_, CyFPhB, and CyTPFB in cell media. All three solutions were measured daily for 5 days and again at 8, 15, and 22 days.

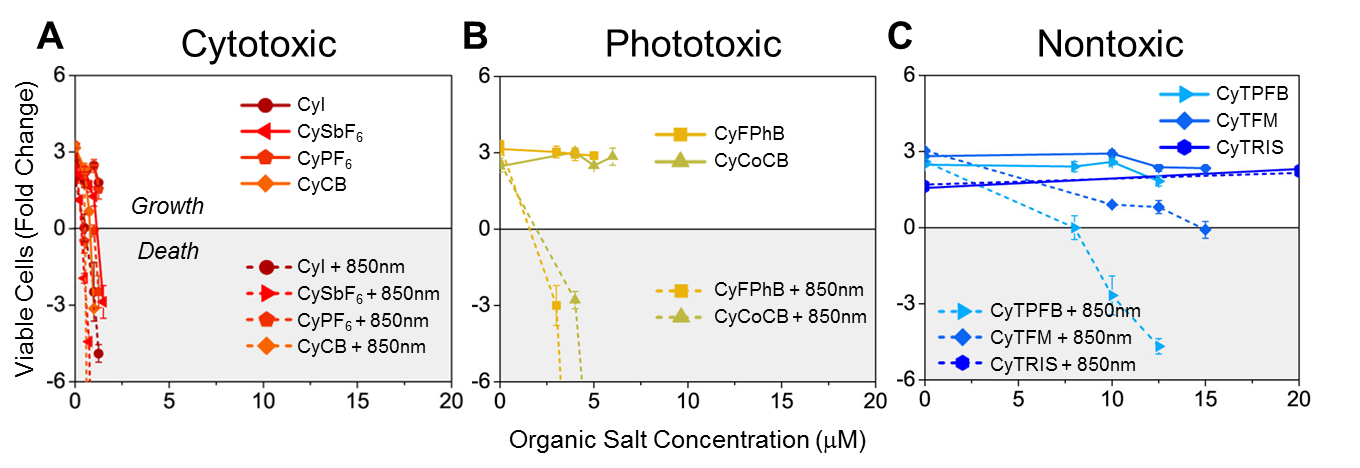

**Supplementary Figure S7.** Metastatic human melanoma WM1158 cells were incubated with various concentrations of Cy+ with different anionic pairings with or without NIR (850 nm) excitation. Cell viability was determined on day 4 by trypan blue staining and cell counting. (**A**) CyI, CySbF_6_, CyPF_6_, and CyCB (red/orange) are toxic at low concentrations (1 μM), and cell death occurs independent of light excitation (cytotoxic). (**B**) CyFPhB and CyCoCB (yellow/green) do not display significant toxicity without light activation, but when photoexcited they induce significant cell death (phototoxic). (**C**) CyTPFB, CyTFM, and CyTRIS (blue) display low toxicity with and without light (nontoxic). This data agrees with the trend observed in A549 cell toxicity. Data are displayed as means ± SEM, *n* = 3

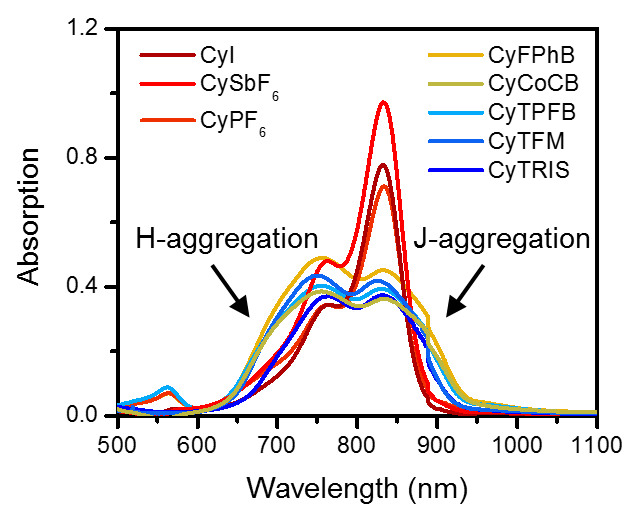

**Supplementary Figure S8.** Salt **n**anoparticle formation in cell media. Organic salts fully dissolved in DMSO have a clear maxima at 830 nm with a leading shoulder when characterized with UV-vis spectroscopy. After nanoparticle formation and introduction into cell media, combinations of H- and J-aggregation of organic salts can still be seen by blue-shifted peaks (lower wavelength) and red-shifted peaks (higher wavelength), respectively, for all but the smaller anions (I^-^, SBF_6_^-^, PF_6_^-^).

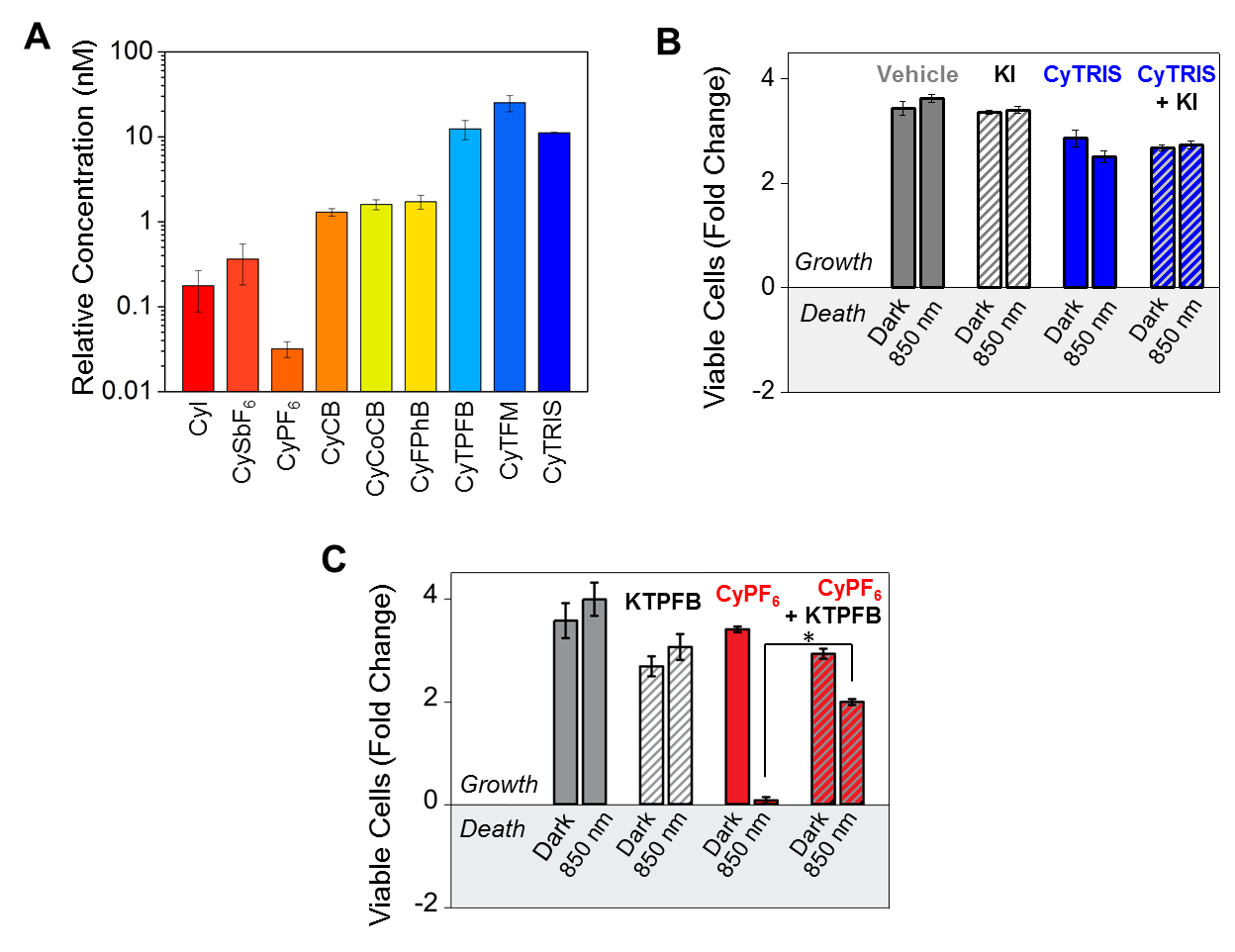

**Supplementary Figure S9**. Tunability in phototoxicity is not due to cellular accumulation or counterion toxicity. (**A**) Intracellular organic salt accumulation by A549 cells was determined using UPLC-MS. In all cases, cells were incubated with 1 μM of indicated organic salt for 30 hours. Data are displayed as means ± SD, *n* = 3. (**B**) Iodide (I^-^) is not toxic when paired with potassium (K^+^), and KI addition does not make CyTRIS toxic. A549 cells were incubated with vehicle, 1 μM KI, 30 μM CyTRIS, or 1 μM KI + 30 μM CyTRIS with or without NIR (850 nm) excitation. Cell viability determined by trypan blue staining and cell counting. Data are displayed as means ± SEM, *n* = 3. (**C**) The phototoxicity and cytotoxicity of CyPF_6_ can be mitigated by the addition of KTPFB, which is not found to be toxic. A549 cells were incubated with vehicle, 15 μM KTPFB, 1 μM CyPF_6_, or 15 μM KTPFB + 0.5 μM CyPF_6_ with or without NIR (850 nm) excitation (**P* ≤ 0.05). Data are displayed as means ± SEM, *n* = 3.

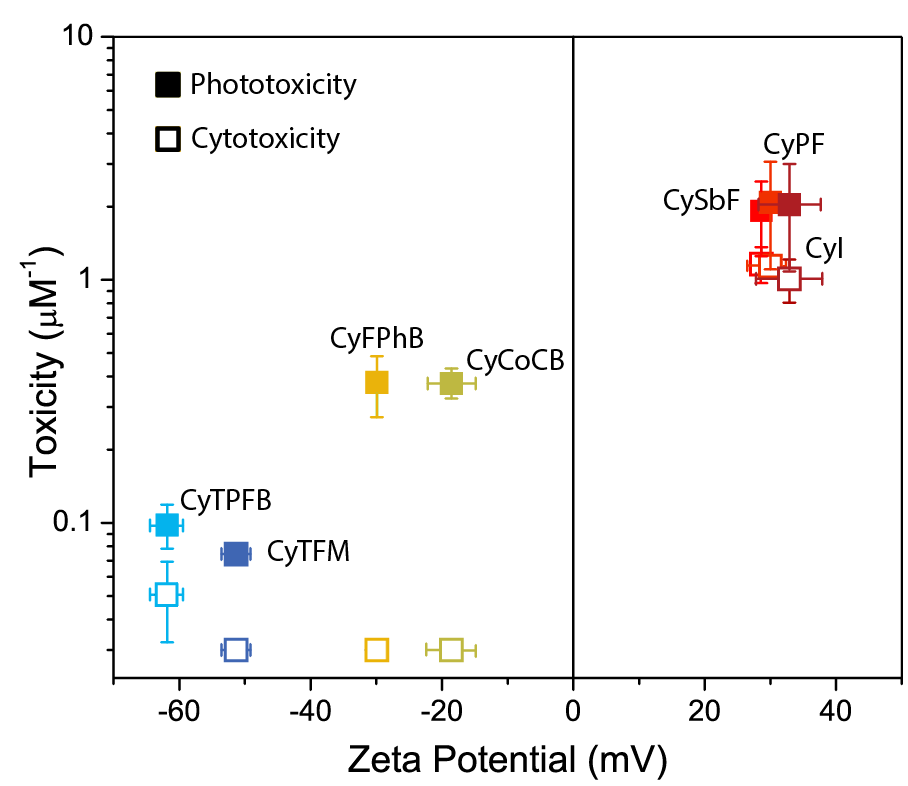

**Supplementary Figure S10.** Cytotoxic anion pairings are found to have positive zeta potentials, while non-cytotoxic pairings display negative zeta potentials. Zeta potential values are provided in Supplementary Table S2. Toxicity values are the inverse of IC_50_ values provided Supplementary Table S4. Data are displayed as means ± SD, *n* = 3.

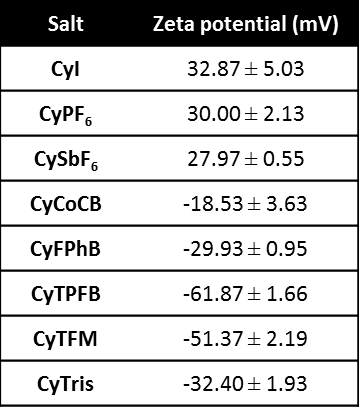
**Supplementary Table S1. Zeta potentials of nanoparticles.** Zeta potential was calculated from electrophoretic mobility using a Malvern Zetasizer NS of organic salt nanoparticles.

**Supplementary Table S2. Quantum yields for CyX and Cy7X salts.**

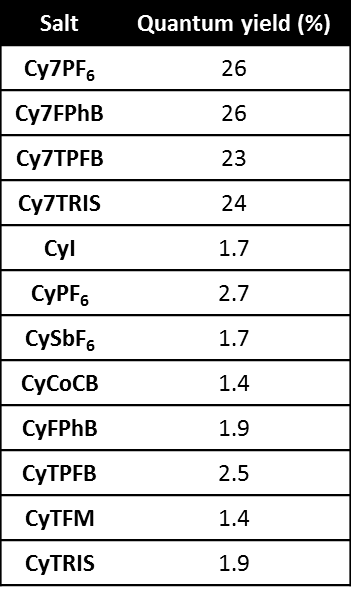

**Supplementary Table S3. Cyanine monomer and nanoparticle solubilities.**

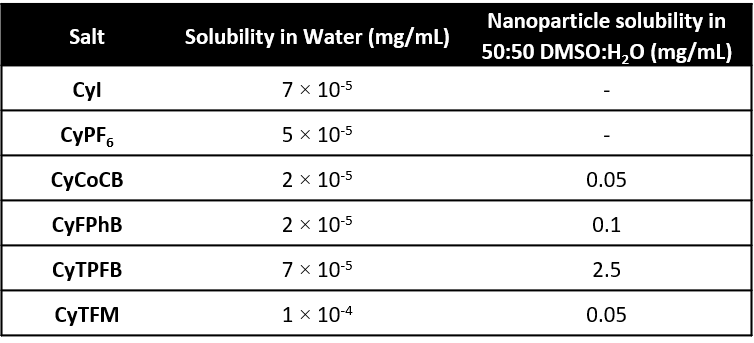

**Supplementary Table S4. Cytotoxicity and phototoxicity of organic salts.** The half maximal inhibitory concentration (IC_50_) values were generated by linear regression analysis for A549 cells. The error is displayed as a 95% confidence interval. NA implies no observable toxicity trend.

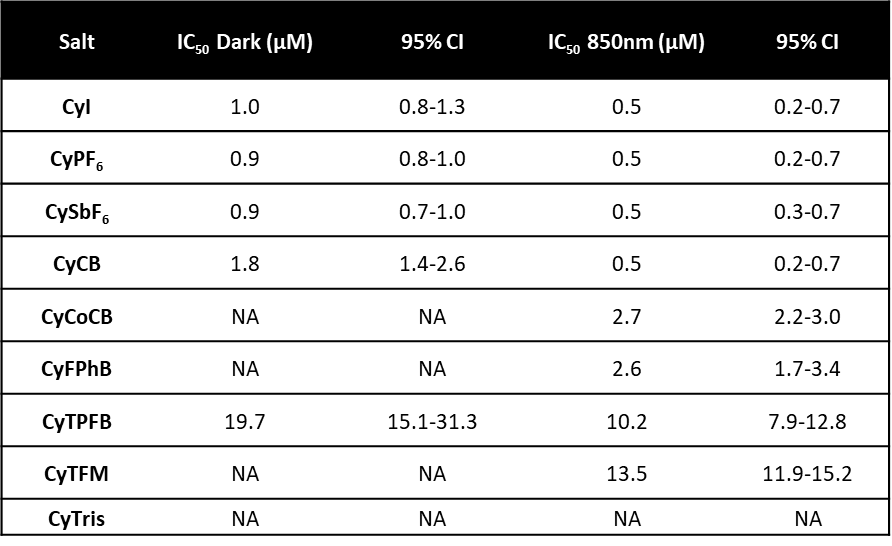

**Supplementary Table S5.** Intracellular localization of the fluorescent ion in A549 cells does not change with counterion. Variables of colocalization that measure the linear relationship between red (organic salt analog) and green (Rhodamine123) fluorescence (Pearson’s coefficient), overlap of red to green area (Mander’s coefficient 1), and overlap of green to red area (Mander’s coefficient 2). All organic salts show a positive linear correlation with mitochondrial fluorescence, with similar degrees of colocalization with mitochondria.

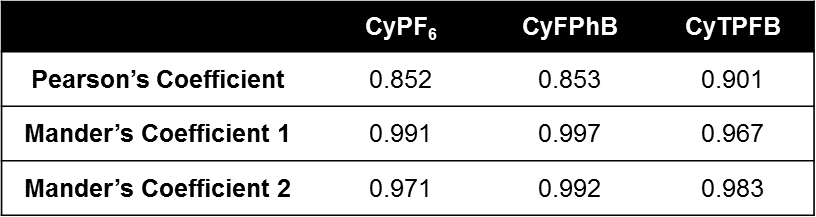
